# Paired sequencing of IgA-bound bacteria reveals widespread associations between adaptive immunity and gut microbiome gene expression

**DOI:** 10.64898/2026.10.05.756684

**Authors:** Logan A. Sauers, Kirsten A. L. Y. Lim, Xiaotong Zhang, Claire Morrisroe, Michal Smiga, Matthew R. Hepworth, Katharine Z. Coyte

## Abstract

Immunoglobulin A (IgA) is the most abundant antibody within the mammalian gut, where it binds to both pathogenic and commensal microbes. Despite its prevalence, the impact of IgA upon the biology and behavior of individual microbes remains poorly understood. Here we present Paired Immunoglobulin Sequencing (PIg-seq), a high-throughput method to identify IgA-associated changes in bacterial gene expression. Using PIg-seq we uncover widespread – and in many cases previously unrecognized – transcriptional responses associated with IgA-binding within the mouse gut. While many of these associations are conserved across diverse bacterial lineages, others vary dramatically even between closely related strains, revealing striking strain-level heterogeneity. Most notably, IgA binding is associated with altered expression of bacterial conjugation genes, and multiple prophages, suggesting an exciting link between adaptive immunity and horizontal gene transfer. Together, our findings establish IgA as a key regulator of microbial behavior, with potential consequences for both the ecology and the evolution of the gut microbiome.

## Main text

Humans and other mammals harbor vast communities of microbes that are essential for host health^1–4^. Realizing these health benefits, however, requires hosts to carefully regulate both the composition and activity of their resident microbes. To achieve this, mammals have evolved a diverse suite of microbiome control mechanisms that promote beneficial taxa while limiting organisms and behaviors that threaten host fitness, among which the immune system is perhaps the most versatile and the most sophisticated^5–8^. However, despite our growing recognition that immunity influences microbial community composition, we still know remarkably little about how the immune system alters bacterial physiology within complex communities.

One fascinating immune mechanism by which hosts may control their microbial communities is secretory Immunoglobulin A (IgA). IgA is the most abundant antibody within the mammalian gut, with approximately 3-5g of IgA secreted into the human gut each day^9^. IgA binds a large proportion of the bacteria within the microbiome, targeting both pathogenic and commensal microbes, potentially in a strain dependent manner^10^. As a consequence, IgA is often thought to play a key role in shaping the composition of the mammalian gut microbiome^11–19^. Indeed, both humans and mice that lack IgA do show detectable differences in gut microbiome composition, as compared to immunocompetent controls^13,14,20^. However, these compositional differences are often modest at best, raising an apparent disconnect between the extensive investment made by hosts into IgA production, and its seemingly limited biological impact. One possible explanation for this disconnect is that a major function of IgA lies not only in determining which bacteria are present, but also in altering the behavior of the bacteria it binds.

Consistent with this hypothesis, there is growing evidence that IgA can directly modulate bacterial physiology. Elegant *in vitro* studies have shown that *Escherichia coli* exposed to IgA downregulate expression of flagellin genes^21^, thus constraining a virulence factor required for penetrating the gut epithelium. This observation has been further supported by *in vivo* observations of elevated flagellin levels in *Rag1*^-/-^ mice, which lack an adaptive immune system^21^. Similar work with the common gut commensal *Bacteroides thetaiotaomicron*^22^ has shown that IgA exposure can downregulate the expression of some polysaccharide utilization (PUL) genes, whilst upregulating the expression of others, in the latter case modulating how *B. thetaiotaomicron* interacts with other members of the gut community^21^. Together, these exemplar studies provide initial evidence that IgA binding can indeed directly reprogramme aspects of bacterial transcription, however, the wider impact of IgA on microbial biology remains poorly understood.

To date most studies have focused on singular, pre-selected bacterial species or pathways, frequently relying on *in vitro* or gnotobiotic *in vivo* systems^17,21–25^. Therefore, we still do not truly understand whether IgA-driven changes in bacterial gene expression are widespread – with many microbes exhibiting similar changes in expression – nor the full extent to which different genes and functions may be affected. We also have relatively little understanding of whether changes observed in reductive *in vitro* or gnotobiotic settings translate to complex microbial communities present within mammalian mucosal environments. Simply put, our knowledge of the relationship between IgA binding and microbial behavior across microbial phylogenies and within natural systems remains severely limited.

To address this gap, here we present Paired Immunoglobulin Sequencing (PIg-seq), a new method which couples IgA separation of *in vivo* derived microbes with paired metagenomic and metatranscriptomic profiling. Using this approach, we uncover widespread differences in bacterial gene expression associated with IgA binding within the intestinal microbiota of conventional mice. Many of these transcriptional signatures are conserved across diverse bacterial lineages, and include several functions previously identified in targeted studies. Perhaps most importantly, we identify numerous genes and functions not previously linked to IgA binding. These include altered expression of various virulence factors and, most notably, increased expression of genes involved in horizontal gene transfer and bacteriophage activity. Finally, we also observe striking differences in IgA-associated gene expression patterns between closely related strains, suggesting hereto unappreciated heterogeneity in the response of individual microbes to IgA binding. Together, our results provide new insights into the ways in which IgA may shape microbial behaviour. More practically, we provide a powerful new tool for disentangling the mechanisms by which IgA may determine microbiome dynamics and consequently host health.

## Results

### Developing Paired IgA sequencing (PIg-Seq)

One approach to understand, at scale, whether IgA influences bacterial behavior is to compare community-wide gene expression profiles between bacteria that are bound by IgA (the IgA-positive fraction) and those that are not (the IgA-negative fraction) within the same microbiome sample. However, differences in RNA abundances between these fractions can arise for two fundamentally different reasons. One possibility is that individual bacterial taxa genuinely express a different suite of genes when they are bound by IgA, as compared to when they are unbound. Alternatively, IgA-positive and IgA-negative fractions may differ in their underlying community composition, such that genes from taxa that are enriched in one fraction appear differentially expressed, simply because these genes are more abundant. To address this challenge, we adapted previously reported IgA-Seq methodologies^10,26–28^ to develop PIg-Seq. This method simultaneously profiles both the metagenome and the metatranscriptome of IgA-positive and IgA-negative bacterial fractions within the same microbiome sample (**Fig 1a**). This paired approach is key, because incorporating metagenomic sequencing data as covariates in differential expression models enables us to disentangle genuine IgA-associated differences in gene expression from those driven solely by differences in the underlying microbiome composition.

**Figure 1.**
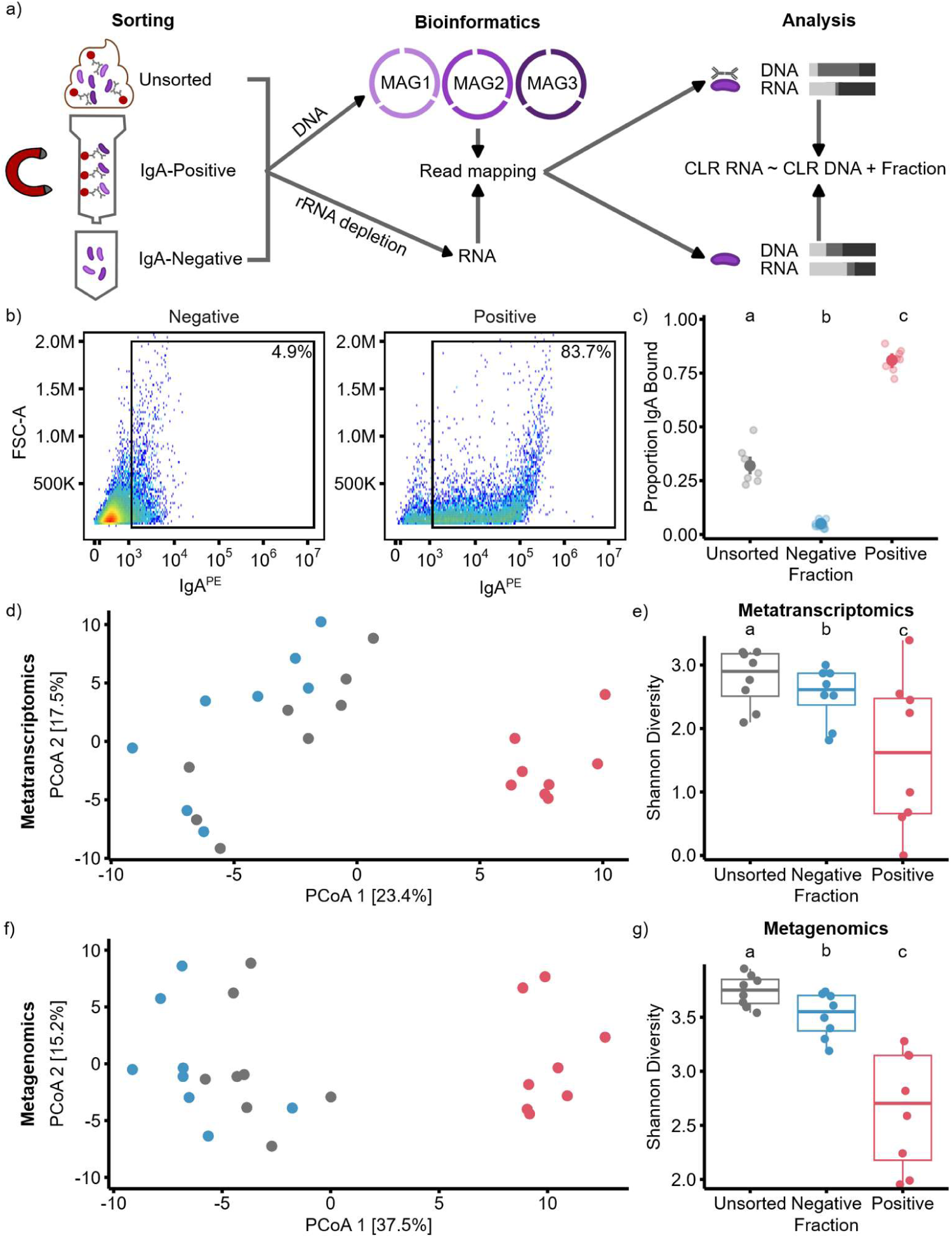
Experimental workflow and characterization of microbial fractions. **a)** Schematic overview of the experimental design, showing separation, bioinformatics, and statistical analysis. **b)** Representative flow-cytometry plots demonstrating separation of IgA-positive and negative bacterial populations. **c)** Proportion of bacteria bound by IgA in the Unsorted, Negative and Positive fractions. Points represent individual samples and point ranges show model-estimated marginal means with 95% confidence. **d)** PCoA of CLR-transformed metatranscriptomic profiles based on Euclidean distances. **e)** Shannon diversity of metatranscriptomic profiles across fractions. **f)** PCoA of CLR-transformed metagenomic profiles based on Euclidean distances. **g)** Shannon diversity of metagenomic profiles across fractions. For panels **d–g**, unsorted, Negative and Positive fractions are indicated by grey, blue and red, respectively.

As a first test, we applied PIg-Seq to fecal samples from 8 adult mice (4 female, and 4 male), generating 24 paired metagenomic and metatranscriptomic profiles. Flow cytometry confirmed efficient separation of IgA-positive and IgA-negative bacteria within each sample (**Fig 1b**), with average IgA-binding rates of 31.93% (+/- 2.11% SE) in the unsorted fraction, enriched to 80.99% (+/- 1.76% SE) in the IgA-positive fraction and depleted to 4.99% (+/- 0.94% SE) in the negative fraction (**Fig 1c and Table S1**). IgA binding was benchmarked against IgMi mice, which lack secretory antibody binding to bacteria, as previously published^29^. We see that IgA-positive fractions clearly separate from both comparison groups in ordination space (F = 3.368, Df = 2, p- value = <0.001, **Fig 1d**). Transcriptional diversity was significantly lower in the IgA-positive fraction compared to either the IgA-negative or the unsorted fractions (F = 7.021, Df = 2, p-value = 0.008, **Fig 1e and Table S2**). However, our metagenomic data also revealed the same overall pattern: IgA-positive fractions clearly separated from unsorted and negative in ordination space (F = 6.4722, Df = 2, p-value < 0.001, **Fig 1f**) and were less taxonomically diverse than either comparison group (F = 24.735, Df = 2, p-value < 0.001, **Fig 1g and Table S3**). This close correspondence between the metatranscriptomic and metagenomic datasets highlights the central challenge of distinguishing changes in bacterial gene expression from changes in underlying community composition.

### IgA-binding is associated with widespread and conserved differences in gene expression ***within* individual organisms**

To determine whether IgA-binding is associated with genuine differences in gene expression, after accounting for underlying differences in gene abundance, we fit a linear mixed effects model on the relative transcriptional abundance of each individual gene within our dataset, using relative gene abundances as covariates^30,31^. This analysis revealed 9,761 genes that were differentially expressed between IgA-positive and IgA-negative fractions, spanning 87 different MAGs from 11 different families (**Fig 2a and Table S4)**. These differentially expressed genes were unevenly distributed, both in number and direction, across bacterial families (**Fig 2b**). For example, members of the *Erysipelotrichaceae* family displayed widespread reductions in activity with IgA-binding, with 37.3% of annotated genes within this family exhibiting significantly lower expression levels in the IgA-positive fractions. In contrast, more than 5% of annotated genes in both *the Rikenellaceae* and *Marinifilaceae* families exhibited higher expression in IgA-positive fractions.

**Figure 2.**
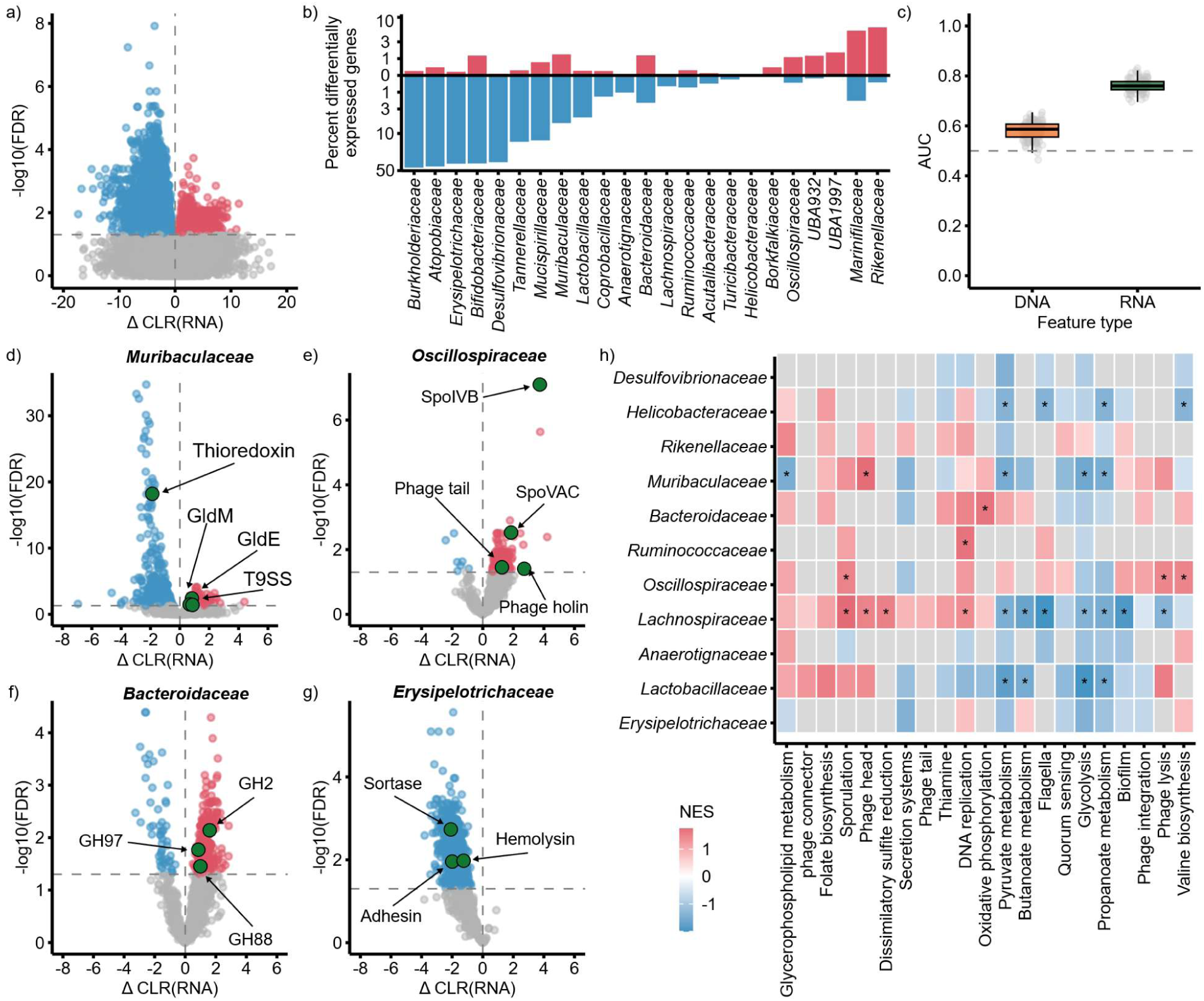
Differential transcriptional activity associated with IgA binding across bacterial genomes and families. **a)** Differential expression analysis of genes within individual metagenome-assembled genomes, comparing the IgA-positive and IgA-negative fractions. **b)** Proportion of genes within each bacterial family classified as significantly upregulated, downregulated, or unchanged. The y-axis is log scaled. **c)** Random forest classifier performance for predicting whether a bin’s relative expression profile originated from the IgA-positive or IgA- negative fraction. The dashed line indicates random classification (AUC = 0.5). **d–g)** Volcano plots showing differential expression of gene products between IgA-positive and IgA-negative fractions for **d)** *Muribaculaceae*, **e)** *Oscillospiraceae*, **f)** *Bacteroidaceae*, and **g)** *Erysipelotrichaceae*. In volcano plots, points are coloured according to statistical significance and direction of differential expression. **h)** Gene set enrichment analysis (GSEA) of functional modules across bacterial families. Gene products were ranked by the differential-expression test statistic, and enrichment is shown as normalized enrichment score (NES); asterisks indicate adjusted *P* < 0.25.

Despite these family specific patterns, however, we were also able to detect evidence for a conserved transcriptional signature of IgA binding across taxa. Specifically, random forest classifiers trained on the relative gene expression profiles of individual MAGs were able to accurately distinguish between IgA-positive and IgA-negative bacterial populations (**Fig 2c**, bootstrapped AUC = 0.761, CI 0.756-0.766). And, crucially, these RNA-based models outperformed equivalent models trained on genome content alone (bootstrapped AUC = 0.580, CI 0.573-0.588). Together, our results suggest that IgA-binding is associated with widespread differences in gene expression *within* individual organisms, and that at least some of these patterns are conserved across diverse bacterial lineages.

### PIg-Seq identifies novel associations between IgA binding and gene expression

We next sought to identify precisely what functions are associated with IgA-binding, initially focusing on functional changes that were shared within families to maximise statistical power. To do so, we combined family-level differential expression analyses with gene set enrichment analyses to identify both individual gene products (**Fig 2d-g and Table S5**) and broader biological processes (**Fig 2h and Table S6**) that were differentially expressed between IgA-positive vs IgA-negative populations. Validating our approach, both independently identified several functional responses that have previously been linked with IgA binding in reductionist systems. These included reduced expression of flagellar synthesis and structure in *Helicobacteraceae* and *Lachnospiraceae* (**Fig 2h**), altered expression of polysaccharide utilization genes in *Bacteroidaceae* (**Fig 2f**), and, in multiple families, reduced thioredoxin expression (**Fig 2f**). The identification of these previously reported functions as being influenced by IgA binding is interesting for two reasons; first, it provides reassurance that PIg-seq can reliably detect genuine associations between IgA-binding and microbial behaviour, even with relatively small sample sizes. But more fundamentally, it suggests a conservation of known IgA-binding responses across multiple, non-model organisms, within real gut environments.

We also identified numerous differentially expressed gene products and pathways that have not previously been associated with IgA-binding (**Fig 2d-h**). In keeping with the IgA’s hypothesized role in suppressing virulence, we found that IgA-positive populations exhibited lower expression of a suite of virulence factors, including sortase, adhesin, hemolysin, and secretion system components (**Fig 2g and Table S5**). IgA-positive populations also exhibited suppression of butanoate, central carbon, and propanoate metabolism, alongside enrichment of sporulation (**Fig 2h**), suggesting a possible shift in IgA-positive bacteria from active growth towards an enrichment of survival responses.

Perhaps most notably, we identified two potential links between IgA-binding and altered patterns of horizontal gene transfer. First, in both the *Muribaculaceae* and the *Bacteroidaceae,* we found that IgA-positive populations exhibited higher expression of genes associated with bacterial conjugation systems (**Fig 3a and Table S7**) – that is, increased expression of the machinery that bacteria use to transfer genetic material to neighboring cells. This included increased expression of genes associated with specific components the relaxosome and Type IV secretion system (e.g. traI, traJ, traK, etc), alongside increased expression of other genes annotated more generically as potentially associated with mobility (e.g. “mobilization protein”, “plasmid transfer protein”).

**Figure 3.**
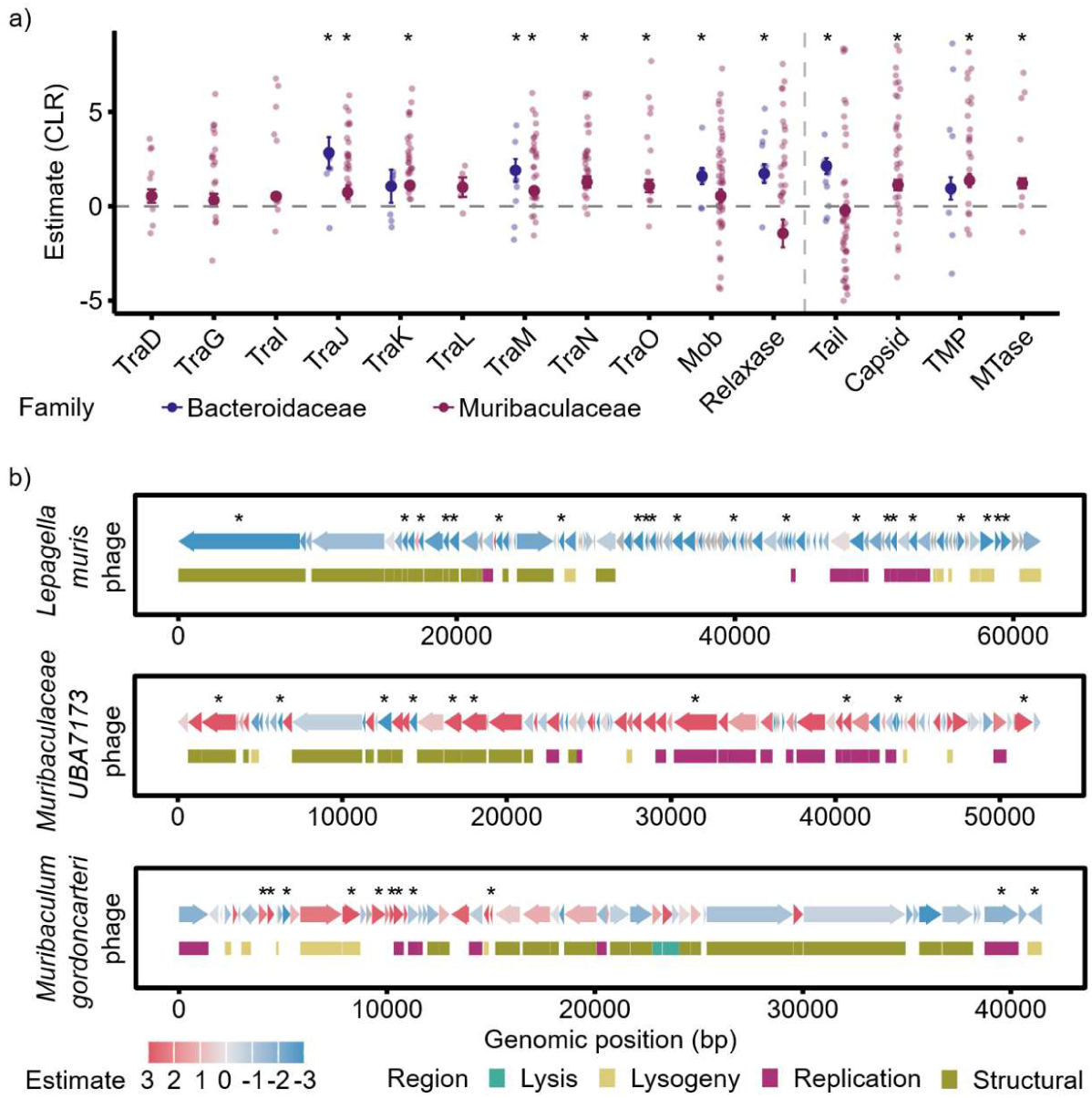
Differential transcriptional activity of horizontal gene transfer systems associated with IgA binding. **a)** Differential expression of conjugative transposon-associated genes, conjugation accessory proteins, and prophage-associated proteins in *Bacteroidaceae* and *Muribaculaceae*. Large points show estimated marginal means, with error bars representing standard errors. Smaller individual points show per-sample, per-bin positive-minus-negative CLR observations. **b**) Genome maps of three phage genomes, showing gene orientation and genomic position. Genes included in the differential-expression analysis are coloured according to their estimated change between IgA-positive and IgA-negative fractions, while genes not tested are shown in grey. Asterisks indicate genes with significant differential expression after multiple- testing correction. Coloured annotation bars indicate predicted gene functional category: lysogeny, replication, structural, and lysis. In all plot asterisks indicate products **(a)** or genes **(b)** with FDR < 0.05.

Secondly, we found that at both the individual gene and the pathway level, IgA-positive populations exhibited strikingly altered expression of genes belonging to prophages – viruses that have integrated into individual bacterial genomes (**Fig 2h**). To test this second observation further, we developed a custom pipeline to identify complete phage genomes and then quantify expression changes within them. This then enabled us to determine whether individual prophages were consistently expressing multiple genes in the same direction – indicating genuine IgA-associated changes in prophage activity (rather than simply noise or sequencing artifacts). This analysis identified multiple prophages that exhibited consistent IgA-associated differences in gene expression across their genomes (**Fig 3b and Table S8**). However, the direction of these differences varied between phages. For example, we identified one phage (found within *Lepagella muris*) which showed consistently lower activity in IgA-positive populations, potentially suggesting suppression of this phage by IgA binding. In contrast, in *Muribaculaceae UBA7173* and *Muribaculum gordoncartori* we found two phages that appeared consistently upregulated in IgA-positive fractions. Closer inspection of these phages revealed that increases in the *Muribaculaceae UBA7173* phage occurred in multiple structural and replication genes, consistent with IgA driving phage induction, whereas in the *Muribaculum gordoncartori* phage these changes occurred predominantly in lysogeny related genes, potentially showing suppression of induction. Together, these findings hint at exciting and previously unrecognized associations between IgA binding and multiple mechanisms of horizontal gene transfer, suggesting that host antibodies may directly influence bacterial evolution within the mammalian gut.

### Closely related taxa can exhibit strikingly different IgA-binding associations

Our initial analyses focused on IgA-associated expression patterns that were conserved within and across bacterial taxa. However, previous work has shown that even closely related strains can differ markedly in their likelihood of being bound by IgA^10^, leading us to wonder whether IgA-associated transcriptional responses might also vary at the strain level. We therefore more deeply investigated our differential gene expression and gene set enrichment analyses at the level of individual MAGs^10^.

These analyses revealed substantial heterogeneity in IgA-associated gene expression patterns across closely related taxa, including in some cases between conspecific strains (**Fig 4a and Table S9**). That is, we observed multiple instances where the same gene or pathway appeared to be enriched in IgA-positive populations for one MAG, yet depleted in IgA-positive populations for another closely related MAG. In some cases, these differences were remarkably widespread, for example we identified two strains of *Muribaculaceae UBA7173* which showed opposing IgA- associations in 28.6% of their shared pathways within the GSEA (**Fig 4b**). Moreover, this heterogeneity also extended to pathways previously associated with IgA binding. For example, bacteria capable of degrading host mucins are thought to be preferentially bound by IgA, however, little is known about whether IgA influences the expression of these genes. We found that one strain of the common gut commensal *Lepegella muris* exhibited consistently higher expression of mucin degradation genes when bound by IgA, yet a second strain exhibited consistently *lower* expression of the same functions (**Fig 4b**) with distinctive divergence between their expression profiles (**Fig 4c**). Together, our results suggest that the interplay between IgA binding and bacterial gene expression can, in some cases, be remarkably strain specific.

**Figure 4.**
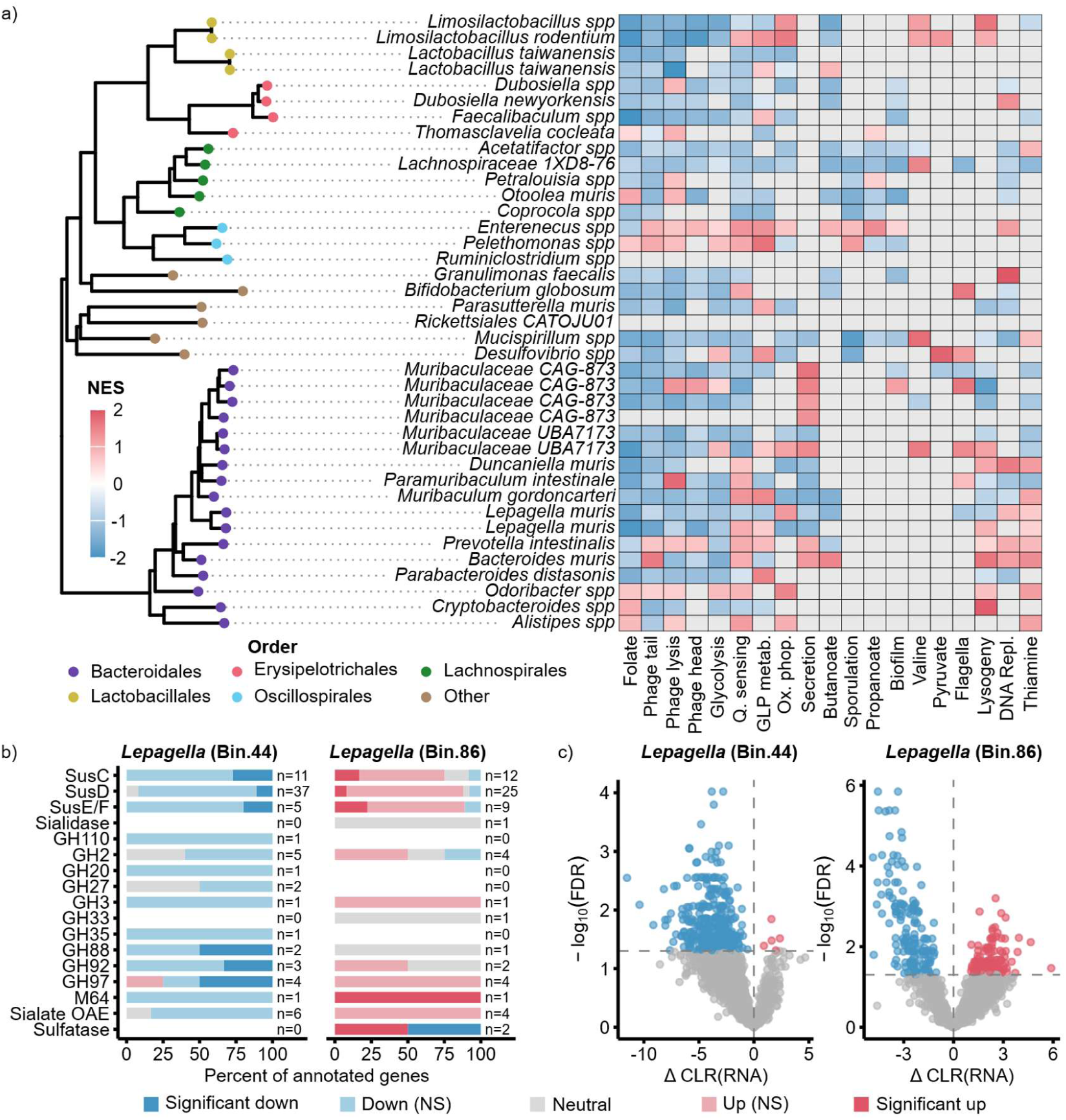
Distinct metabolic responses between closely related strains associated with IgA binding. **a)** Phylogenetic tree of the selected bacterial bins and GSEA heatmap showing normalized enrichment scores (NES). Heatmap columns represent functional modules, ordered according to their correlation in enrichment profiles across bins. **b)** Differences in polysaccharide- utilization gene composition in *Lepagella* bins. Bars show the proportion of annotated genes within each functional category, classified according to the magnitude and significance of differential expression between positive and negative fractions. Numbers next to bars indicate the number of annotated genes. Non-significant genes were classified as directionally up or down if the model estimate exceeded a log2 fold-change threshold of ±log2(1.5) between the Positive and Negative fractions. **c)** Volcano plots showing differential expression of genes in two *Lepagella muris* bins. Points are coloured according to statistical significance and direction of differential expression.

### Gene expression predicts probability of IgA binding

It is well established that IgA is more likely to bind some bacterial taxa than others, however recent work suggests that this binding propensity can vary considerably even between closely related strains^10^. As a final question, we therefore asked whether a given organism’s propensity to be bound by IgA could be predicted from its behavior within the gut. To do so, we calculated the IgA-binding score per sample for each MAG within our dataset (**Fig 5a and Table S10**), then categorised each MAG (per sample) as either “low binding”, “medium binding” or “high binding”. We then trained three independent sets of random forest classifiers to predict binding category: the first set of classifiers used just the genome content of each MAG, the second used the relative gene expression of each MAG when bound by IgA, and the third used the relative gene expression of each MAG when not bound by IgA.

**Figure 5.**
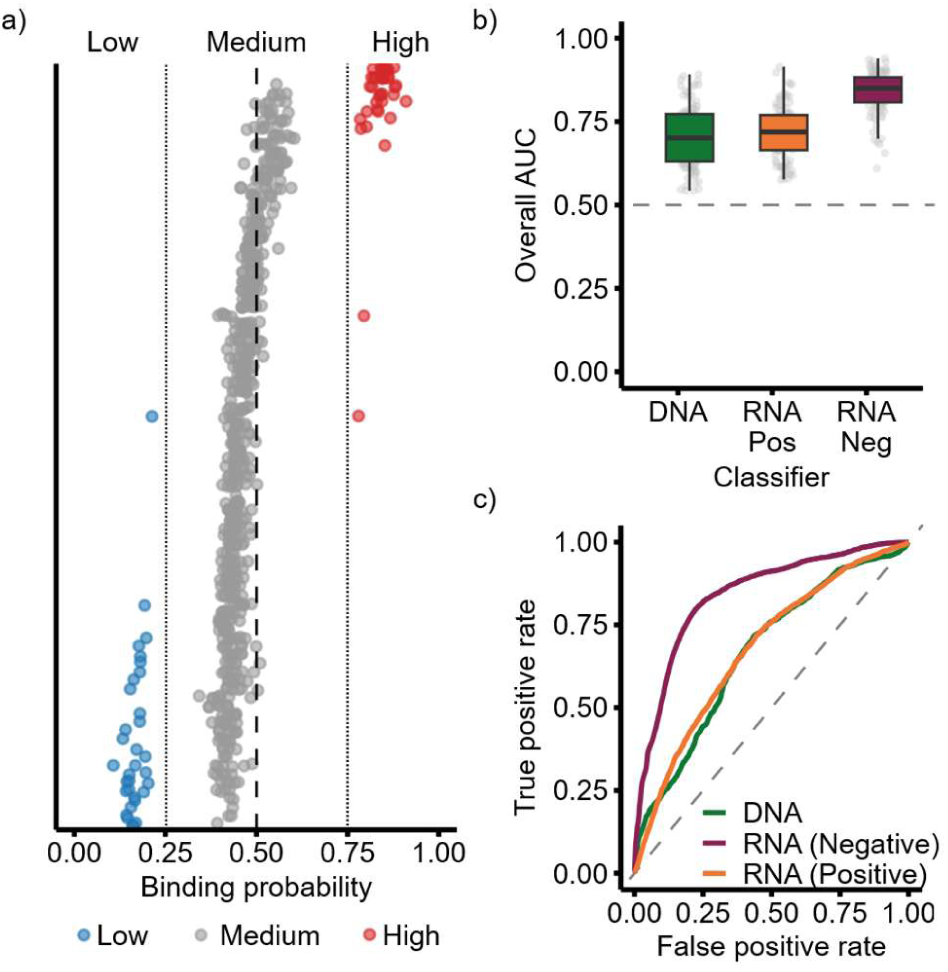
Transcriptional features of IgA-negative bacteria predict IgA binding intensity. **a)** IgA binding scores for taxa. Dots represent the sample determined binding scores of metagenome assembled genomes. Values greater than 0.75 were classified as high binding, values between 0.25 and 0.75 classified as medium binding and those below 0.25 were low binding. **b)** Area- under-the-curve (AUC) distributions for predicting bacterial IgA-binding class from three trained classifiers. Prediction was performed using random-forest models with either DNA presence/absence, negative fraction relative expression profiles, or positive fraction relative expression profiles as input features. Boxes show the distribution of AUC values, and the dashed line indicates random classification (AUC = 0.5). **c)** Receiver operating characteristic curves illustrating classification performance of the genomic models for distinguishing IgA-binding intensity.

Surprisingly, we found that models trained on relative gene expression within the *IgA-negative* fraction were able to predict each MAG’s IgA-binding category with remarkably high accuracy (RNA Negative macro-averaged AUC = 0.838, 95% CI: 0.685-0.929). These models outperformed equivalent models trained on genome content alone (Wilcoxon p = 1.60 × 10^⁻20^), and models trained on gene expression within the IgA-positive fraction (p = 8.02 × 10^⁻21^; **Fig 5b and c**). Notably, the RNA Negative model performed well across all three binding categories, particularly for the minority High-binding class (per-class AUC = 0.907, 95% CI: 0.605-0.996). Our results should be interpreted with some caution, as relatively few MAGs were identified as either preferentially bound, or preferentially unbound, within our dataset. Nonetheless, they raise the interesting possibility that preferential IgA targeting is driven not only by bacterial taxonomy, but also by how bacteria behave within the gut.

## Discussion

Secretory IgA is thought to be a key immune pressure by which mammals control the dynamics of their gastrointestinal microbiome. Yet remarkably little is known about whether and how IgA binding affects bacterial behavior within natural communities. Here we developed a new method – PIg-seq – that enables us to address this gap, systematically identifying associations between IgA binding and bacterial gene expression at scale and within natural microbiomes. The key novelty in our method lies in our pairing of metagenomic and metatranscriptomic sequencing from the same IgA-separated samples, enabling IgA-associated differences in gene expression to be distinguished from underlying differences in bacterial abundances. Applying our approach to the mouse gut reveals that IgA-associated differences in bacterial gene expression are not only widespread across diverse bacterial lineages but also encompass a far more diverse array of functions than previously recognized.

Perhaps our most exciting finding is the observation of repeated associations between IgA binding and altered expression of genes involved in horizontal gene transfer. Across multiple taxa, we found that IgA-positive populations consistently exhibited higher expression of genes associated with conjugation – a major mechanism by which bacterial cells exchange genetic material^32–35^. We also observed altered expression of prophages – viruses that integrate into the bacterial genome and thus can act as vectors for genetic material^36,37^ – although here the responses were more heterogenous. Several mechanisms could be driving these observations. For example, one possibility is that IgA-mediated agglutination of cells drives a local increase in bacterial density^38^, creating conditions that simultaneously promote conjugation and suppress (or trigger) prophage induction^19,39,40^. Alternatively, IgA may preferentially target cells already undergoing horizontal gene transfer, potentially even facilitating their removal from the gut.

While our data cannot distinguish between these and other possibilities, nonetheless they raise the fascinating possibility that IgA could shape not only the ecological, but also the evolutionary dynamics of the gut microbiome.

Our results also add weight to our growing understanding that IgA-microbe interactions can, at least in some cases, be remarkably specific^21^. Consistent with previous studies, we found that closely related strains can differ substantially in the propensity to be bound by IgA. However, what we now show is that closely related strains may also exhibit very different responses to IgA binding, with members of the same species sometimes displaying completely opposing patterns of IgA-associated gene expression. These observations highlight the importance of strain-level resolution for understanding immune–microbiome interactions.

While our results are interesting, several important caveats should be acknowledged. First, thus far we have applied our method to only a modest cohort of healthy mice. Second, more conceptually, while PIg-seq can identify associations between IgA binding and gene expression, it cannot, on its own, identify causal relationships. That is, PIg-seq cannot distinguish whether IgA binding actively induces changes in bacterial transcription, or whether bacterial cells with particular expression profiles are simply more likely to be bound by IgA than others. In practice, both scenarios are likely to be at play. For example, we suspect that our observation of reduced flagella expression in IgA-positive populations reflects a direct transcriptional response to IgA binding, as this phenomenon has previously been observed in *in vitro*^36,37^. In contrast, we suspect that the reduced expression of thioredoxin in IgA-positive populations may instead reflect preferential IgA binding of cells that express lower levels of this IgA-modifying enzyme^42,43^. Indeed, our observation that preferential IgA binding can be accurately predicted from bacterial gene expression profiles in the absence of IgA, suggests that IgA may often target bacteria based not only on who they are, but also on what they are doing.

Despite these caveats, PIg-seq provides an advance on existing methodologies for connecting host immunity with microbial function directly within complex communities. It is readily applicable to diverse microbiomes and disease settings in which IgA may implicated, including inflammatory bowel disease, necrotising enterocolitis, and early-life microbiome development. More broadly, our findings suggest that IgA should be viewed not just as a mechanism for regulating microbial composition, but also as a key regulator of microbial physiology, ecological interactions, and horizontal gene transfer. Understanding these processes will be essential for determining how adaptive immunity shapes the function and evolution of the gut microbiome.

## Materials and Methods

### Sample collection

Samples were collected from heterozygous IgMi mice established in the University of Manchester Biological Services Facility. The mice established were originally a gift from A. Waisman (Institute of Molecular Biology, Mainz)^29^. Mice were cohoused with liter mates of mixed genotypes with *ad libitum* feeding and 12 hour day night cycles. Fecal pellets were collected in pre-weighed tubes and flash frozen by placing on dry ice. Tubes were then weighed again with fecal pellet to determine the weight of the pellet. Samples were collected in two batches (**Table S11**).

### Fecal pellet pre-processing

Fecal pellets were thawed on ice and then disrupted by adding 450 µL of chilled PBS and smashing the pellet with pipette tips. A further 450 µL of chilled PBS was then added and pellets briefly by pulsed (3-5 second) vortex to mix until homogenized. Fecal homogenates were then centrifuged at 1,200 rpm (∼150 x G) for 5 minutes in a microcentrifuge prechilled to 4°C. After centrifugation supernatant was removed and transferred to a clean microcentrifuge tube (the initial pellet should mostly contain food debris and was therefore discarded^29^). The supernatant was then centrifuged at 10,000 x G for 5 minutes in a prechilled microcentrifuge with the resulting pellet used for magnetic activated cell sorting (MACs).

### Magnetic activated cell sorting

Bacterial pellets were resuspended in 500 µL of FACs buffer (1mM EDTA and 2% FBS). 100 µL (20%) of this resuspension – representing the “unsorted” cell fraction for DNA/RNA extraction – was transferred to a clean microcentrifuge tube containing 400 µL of FACS buffer and then stored at 4°C until DNA/RNA extraction the same day. The remaining 400 µL (80%) of the resuspension was centrifuged at 8,000 x G for 5 minutes in a prechilled microcentrifuge.

Supernatants were discarded, pellets resuspended in 100 µL of anti-mouse IgA:PE conjugated antibody (1µg/mL) stain and incubated at 4°C for thirty minutes. After incubation samples were washed with 1,000 µL of FACS buffer and pelleted. The bacterial pellets were then resuspended in 1,000 µL of FACS buffer and 100 µL (10%) stored at 4°C to serve as the unsorted bacterial population for flow cytometry. 100 µL of anti-PE beads (Miltenyi Biotec) were added to the remaining 900 µL of sample and samples incubated at 4°C for thirty minutes. After incubation samples were washed in 1,000 µL of FACS and pelleted. Pellets were resuspended in 1,000 µL of FACS and subsequently transferred into a 15mL falcon tube. Microcentrifuge tubes were washed 4 more times with 1,000 µL of FACS with each wash transferred to the same falcon tube as the original pellet. Magnetic cell separation was done using LS Columns (Miltenyi Biotec) and QuadroMACS separator (Miltenyi Biotec). Columns were activated on the magnet by adding 3mL of FACS buffer with discarded flow. To elute the IgA negative (depleted) bacterial population, samples were then added to the columns 2.5mL at a time and flow through collected in falcon tubes. Columns were washed with 3mL of FACS and flow through collected in the same tube. This wash step was repeated once. Columns were removed from the separator and placed in new 15 mL falcon tubes. To elute the IgA positive (enriched) bacterial populations 5mL of FACS buffer was added to the tube and forcefully expelled using the provided plunger. The separation was repeated a second time on the IgA positive population to increase the purity of the enrichment. Falcon tubes containing the IgA negative and positive populations were then centrifuged at 4,500 rpm for 10 minutes in a prechilled centrifuge with soft breaking enabled.

Supernatant was removed and pellets resuspended in 1,000 µL of FACS buffer. 200 µL of these populations were transferred to a clean microcentrifuge tube and topped to 1mL for flow cytometry. The remaining 800 µL saved at 4°C for DNA/RNA extraction the same day.

### Flow cytometry

Controls for the flow cytometry were made from the unsorted bacterial population aliquots. 10 µL (10%) from these 100 µL aliquots were mixed in a clean microcentrifuge tube and 990µL of FACS buffer added. 333 µL were then transferred to three clean microcentrifuge tubes for unstained, SYTO9 (Fisher Scientific), PE controls. Control and sample aliquots tubes for unsorted, bacterial populations were all topped to 1mL of FACS. Controls and all flow aliquot, including unsorted, negative, and positive bacterial population aliquots, were then spun at 8,000 x G for 5 minutes in a prechilled microcentrifuge. Supernatant were discarded and pellets resuspended in 100 µL of SYTO9 stain (8.3µM) and incubated at 4°C for thirty minutes.

Samples were topped to 1mL and washed by gentle inversion followed by centrifugation to pellet. Supernatant was removed, samples resuspended in 200 µL of FACS buffer and loaded onto a 96 well plate for flow. Flow was done using a CytoFlex S Cytometer (Beckman Coulter) with a flow rate of 30µL/minute with the following gains: forward scatter 200, side scatter 100, FITC channel 50, PE channel 200, and a threshold set at 50,000 for forward scatter. Samples were run for 100 seconds, sampling a total of 50 µL with 15 µL of water run between each sample to clean the flow cell.

### DNA/RNA extraction

DNA/RNA extraction was done using the ZymoBIOMICs DNA/RNR Miniprep Kit (Zymo Research). The 800 µL aliqouts saved for DNA extraction from the unsorted, negative, and positive bacterial populations were spun at 8,000 x G for 5 minutes in prechilled microcentrifuge to pellet cells. Pellets were resuspended in 750 µL of DNA/RNA Shield (Zymo Research) and vortexed at 1,800 rpm for 20 minutes in a Fisherbrand Digital Vortex Mixer (Catalogue #15796959) with 24 microtube adaptor (Catalogue #15766959). After lysis extractions follow the kit provided instructions for the separate DNA/RNA extraction with elution being down twice with 25 µL in each elution step for a final eluted volume of 50 µL.

### DNA/RNA sequencing

DNA/RNA sequencing was done in two sets, corresponding to the sets of collected samples. The first set was sent to Novogene (UK) Company Limited. DNA and RNA concentrations of the first set samples were assessed with a bioanalyzer by Novogene and subsequently eukaryotic and prokaryotic rRNA removed and libraries prepared by Novogene following their internal protocols and quality assessed with bioanalyzer and qPCR before sequencing. Sequencing was done on the NovaSeq X Plus Series (PE150) at 12 G of raw data per cell population. The second set of samples were rRNA depleted using the NEBNext Bacterial rRNA Depletion Kit (New England Biolabs #E7850L). Depleted RNA and genomic DNA was submitted to the University of Manchester Genomic Technologies Core Facility (GTCF) and libraries were generated using the Illumina® Stranded mRNA Prep Ligation kit (Illumina, Inc.) and and Illumina DNA Prep kit (Illumina, Inc.). The depleted mRNA was fragmented under elevated temperature and then reverse transcribed into first strand cDNA using random hexamer primers and in the presence of Actinomycin D to improve strand specificity whilst mitigating spurious DNA-dependent synthesis. Following removal of the template RNA, second strand cDNA was then synthesized to yield blunt-ended, double-stranded cDNA fragments. Strand specificity was maintained by the incorporation of deoxyuridine triphosphate (dUTP) in place of dTTP to quench the second strand during subsequent amplification. Following a single adenine (A) base addition, adapters with a corresponding, complementary thymine (T) overhang were ligated to the cDNA fragments. Pre-index anchors were then ligated to the ends of the double-stranded cDNA fragments to prepare them for dual indexing. A subsequent PCR amplification was then used to add the index adapter sequences to create the final cDNA library. DNA prep was done following the standard manufacturer protocol. Library quality was assessed using 4200 Tapestation (Agilent Technologies) with a D1000 ScreenTape and molarity checked with qPCR using KAPA Library Quantification Kit (F. Hoffman-La Roche Ltd). The adapter indices enabled the multiplexing of the libraries, which were pooled prior to loading on to the appropriate flow- cell. Pooled libraries were then paired end sequenced (59 + 59 cycles, plus indices) on an Illumina NovaSeq6000 instrument using the SP lane. Finally, the output data (**Table S12**) was demultiplexed and BCL-to-Fastq conversion performed using Illumina’s bcl2fastq software, (version 2.20.0.422).

### Bacterial bioinformatics

Raw sequencing data was assessed for initial quality using FastQC (version 0.11.5) and then run through a custom cleaning pipeline consisting of removing poor quality reads with Trimmomatic (version 0.39, minlen 36, leading 3, trailing 3, sliding window 4:20)^44^ and host reads removed with bowtie (version 2.3.5.1)^45^ set to very sensitive using the mouse reference genome available on NCBI (GRCm38) (**Table S12**). Cleaned reads from the metagenomic sequencing were assembled using metaspades (version 3.15.5)^46^ within the metaWRAP wrapper (version 1.3.2)^47^. Metagenomic assemblies were binned into genomic units using metabat2 (version 2.12.1)^48^ and maxbin2 (version 2.2.4)^49^ inside the metaWrap wrapper and with concoct (version 1.1.0)^50^ using customized scripts and environments. Bins were refined using the metaWRAP’s updated binning_refiner.py^51^ using a completion threshold of 90% and a contamination threshold of 5% assessed by CheckM1 (version 1.0.18)^52^ within metaWRAP. After refinement, bin quality and completion was reassessed using CheckM2 (version 1.1.0)^53^ and then dereplicated to a non-redundant bin set (**Table S13**) at 0.99 average nucleotide identity with drep2 (version 3.5.0)^54^. Dereplicated bins were then annotated using bakta (version 1.11.4)^55^ and classified using GTDB-TK (version 2.5.2) with database release 226 (**Table S13**)^56–58^. Cleaned RNA and DNA reads were then mapped back to the dereplicated bin set using bwa-mem2 (version 2.2.1)^59^. Reads with map scores below Q20 were excluded from subsequent analysis. Gene counts were obtained using featureCounts (version 2.0.6)^60^ from the subread package^61^ set on paired-end mode and discard multiply mapped reads. Custom bash scripts were used to merge the annotation files, normalize the count tables to TPM for RNA reads and TPM equivalent for DNA reads, and merging these final output files. CoverM (version 0.7.0)^62^ was used to calculate the TPM of each bin for both the DNA and RNA sequencing to measure bin level abundance and transcriptional activity for each bin.

### Phage bioinformatics

Phage genomes were predicted from the dereplicated bin set using geNomad (v1.11.2)^63^. The resulting phage genomes were then filtered by completeness. Retaining only those classified as medium-high quality by CheckV with no quality warning flags^64^. As some host bacteria carried multiple phage genomes, these were dereplicated at 95% average nucleotide identity with ≥85% target coverage using all-vs-all BLASTn, following established species-level thresholds for viral genomes^65^. The resulting non-redundant phage genomes were annotated using Pharokka^66^, phold^67^, and Phynteny^68^, with functional annotations merged across tools^69^ (**table S14**). RNA reads were then mapped back to these genomes as previously described and filtered to MAPQ > 20.

### Statistical analysis

Data analysis was done in R (version 4.3.2)^70^, RStudio (version 2025.9.2.418)^71^, and Jupytner Notebook (version 7.5.5)^72^. The following packages were generally applied throughout the analyses: tidyverse^73^ to organize data frames and structure, furrr, future, and progressr to handle parallelizing the workflows, ggplot2^74^ to create graphics.

### IgA binding enrichment

The enrichment of IgA binding was modelled by fitting a beta distribution linear model, ideal for data in the range 0 to 1, to the proportion of IgA bound bacteria for each fraction using the glmmTMB package^75^. Marginalized estimated means were calculated with the emmeans^76^ package and FDR correction used to control multiple comparisons between fractions.

### Community composition and diversity

Bin abundance from both the RNA and DNA data was analyzed using the phyloseq package. First, bin abundance estimates from CoverM were CLR-transformed. Euclidean distance was calculated and vegan^77,78^ used to perform PERMAnova with 10,000 permutations. Ordinations were plotted with phyloseq and ggplot2^74^. Shannon alpha diversity was calculated for each fraction using phyloseq. Differences in Shannon diversity were assessed using linear mixed effects models with Fraction as a fixed effect and Mouse as a random intercept, followed by pairwise contrasts with FDR correction.

### Phylogenetic tree construction

The phylogenetic tree of the dereplicated bin set was generated with GTDB-TK^57^ in Newick format. The tree was imported R with the package ape^79^. Bins corresponding to those used in the GSEA analysis were selected and analyzed together with the *Chlamydia* bin as the outgroup. Taxonomic information was mapped onto tree tips with tips labelled by species and colored according to Order. The tree visualization was made using ggtree ^80^.

### Differential gene expression analysis

Genes differentially expressed between fractions were identified using linear mixed effects models on CLR-transformed RNA TPM values, with CLR-transformed DNA TPM included as a covariate to account for compositional shifts in the underlying community. The accuracy of linear models with DNA covariates for metatranscriptomic differential expression is supported in previous literature^30,81^. Two levels of analysis were conducted. First, at the bin level, individual genes were modelled with the formula CLR(RNA) ∼ CLR(DNA) + Fraction + (1|Mouse), with FDR correction applied per contrast within each bin. Second, at the family level, TPM values from genes sharing the same product annotation within a family were summed before CLR transformation and modelled with CLR(RNA) ∼ CLR(DNA) + Fraction + (1|Mouse) + (1|Bin), where the additional random intercept for Bin accounts for multiple MAGs contributing to each family. Family-level products were required to be present in at least 2 bins. For both levels of analysis, genes were filtered to require: i) non-zero DNA and RNA TPM values, ii) non-zero variance in both DNA and RNA TPM, iii) no annotation as ribosomal RNA or hypothetical protein, and (iv) at least 4 observations per fraction. FDR correction was used to control for multiple tests.

### Gene set enrichment analysis

Gene set enrichment analysis was performed using fgsea to identify broader functional processes differentially expressed between IgA-positive and IgA-negative fractions. Custom gene sets were constructed by querying gene annotations using regular expression matching (regex) guided by the Kyoto Encyclopedia of Genes and Genomes (KEGG)^82^ and the Prokaryotic Virus Remote Homologous Groups database (PHROGs)^83^. A two-layer exclusion approach was used to minimize false positive rates. First a standard exclusion for hypothetical proteins, domain-containing proteins, and putative annotations was applied. Second gene-set-specific exclusions were identified through manual curation of the list against reference databases. This keyword-based approach was required because Bakta annotation does not prioritize database identifiers, with a median of 72.7% (IQR 65%–77.9%) of genes in KEGG-derived gene sets missing KO number annotations despite having proper functional annotations. The resulting coverage of each custom gene set against its reference database (**Table S15**) had a median of 92.3% (IQR 75.5%–95.9%). All exclusion and regex patterns are available in the publicly available analytical code.

GSEA^84,85^ was conducted at two levels. At the bin level, genes were ranked by t-ratio from the differential expression models. GSEA was attempted only for bins with at least 100 modelled genes, with a minimum of 5 genes required per gene set within a bin. At the family level, products were ranked by t-ratio from the family-level models, with a minimum gene set size of 5. Gene sets were considered significant at the exploratory GSEA threshold of FDR-adjusted p < 0.25, with those reaching FDR-adjusted p < 0.05 considered nominally significant^84,85^.

### Phage differential gene expression analysis

Phage genes differentially expressed between fractions were identified using linear mixed effects models on CLR-transformed RNA TPM values. Unlike the bacterial gene expression analysis, CLR-transformed DNA abundance of the host bacterium (at the bin level) was used as a covariate to control for variation in the host bacterial abundance across fractions. This approach is conceptually analogous to PropagAtE^86^, which uses prophage-to-host DNA coverage ratios to detect phage activation from metagenomes, but applied here to transcriptional data. The model formula was CLR(RNA) ∼ CLR(host DNA) + Fraction + (1|Mouse), with FDR correction applied per contrast across all phage genes. Phage genes were filtered to require at least 2 observations per fraction, presence in at least 2 fractions, and a minimum of 4 total observations. To assess whether individual prophages exhibited consistent IgA-associated changes in expression across their genomes, pharokka gene coordinates were joined with differential expression results for each provirus, enabling visualization of expression changes across complete prophage genomes.

### Random forest classifications

To test whether IgA-positive and IgA-negative bacterial populations exhibit conserved differences in gene expression, a random forest classifier was trained to predict whether a relative expression profile (REP) was derived from the IgA-positive or IgA-negative fraction. First IgA binding scores were calculated using the IgAScores package^28^. The resulting scores range from −1 to 1, which we rescaled to (0, 1) using (score + 1) / 2. Following this, REPs were calculated for each MAG in each sample by applying rarefaction to equalize sequencing depth between fractions, summing RNA TPM values for each unique gene product, and dividing by the total RNA TPM for that MAG, following the removal of ribosomal genes and hypothetical proteins. To avoid classification being driven by differences in IgA binding rate, only MAGs with equal binding probability (IgA score not significantly different from zero) were included. For each of 100 bootstrap iterations, approximately 20% of MAGs were held out as test data, with all samples from a given MAG assigned to either training or testing. Classifier performance was assessed using AUC-ROC^87^. This procedure was repeated using metagenomic relative abundance profiles (DNA) as a control, and differences in AUC distributions between RNA and DNA classifiers were assessed using Wilcoxon rank-sum test^88^.

To test whether IgA binding could be predicted from microbial gene content or transcriptional activity, three independent sets of random forest classifiers were trained to predict per-sample binding status. IgA binding scores were assigned to each MAG in each mouse and categorized as low (< 0.25), medium (0.25–0.75), or high (> 0.75), allowing the same MAG to carry different binding labels across hosts. The first classifier used genome-level DNA presence/absence profiles, where a gene was scored as present if it was ever detected at the DNA level for that MAG across any mouse or fraction. The second and third classifiers used relative gene expression profiles (TPM per product divided by total TPM per MAG) calculated independently within the IgA-positive and IgA-negative fractions, respectively. All three classifiers shared a common feature set, filtered to gene products detected in at least 10% of observations in both fractions, and hypothetical proteins and ribosomal RNA genes were excluded. For each of 100 bootstrap iterations, approximately 20% of MAGs were held out at the bin level, with rejection sampling to ensure representation of all three classes in each test set. Inverse class-frequency weighting was applied during training to account for class imbalance. Classifier performance was assessed using macro-averaged AUC-ROC^87^, differences between classifiers were assessed using Wilcoxon rank-sum tests^88^ on the 100-seed AUC distributions.

## Acknowledgements

We thank the University of Manchester IT core facilities for computational support, and the University of Manchester BSF staff for assistance with animal husbandry and maintenance. We also thank Luke Roberts for allowing us access to equipment, Stacey Holden and Peter Briggs for support with the transcriptomic and metagenomic sequencing and members of the Coyte lab for valuable input.

## Author contributions

Conceptualization: LS, XZ, MRH, KZC. Investigation: LS, CM, MS, XZ. Data curation: LS. Formal analysis: LS, KL, KZC. Funding acquisition: LS, XZ, MRH, KZC. Supervision: MRH, KZC. Original draft: LS, CM, KL. Review and editing: LS, MRH, KZC.

## Data availability

The sequencing data generated in this study has been deposited in ENA under accession PRJEB127118. Data from the bioinformatics pipelines, used in data analysis and sample metadata are available from FigShare (DOI: https://doi.org/10.48420/33951214). Relevant intermediate data and statistics are available within the paper and its Supplementary Information. Scripts for data analysis are accessible from SauersALogan/PIg-Seq.

